# Pangenome insights into structural variation and functional diversification of barley CCT motif genes

**DOI:** 10.1101/2025.03.19.643958

**Authors:** Zihao Zhu, Nils Stein

## Abstract

CCT motif genes play a key role in barley development and flowering, yet their genetic diversity remains underexplored. Leveraging a barley pangenome (76 genotypes) and pan-transcriptome (subset of 20 genotypes), we examined CCT gene variation and evolutionary dynamics. Motif-based searches, combined with genome assembly validation, revealed annotation limitations and novel frameshift variants (e.g., *HvCO10*), indicating active diversification. Pangenome-wide phylogenetic analysis identified clade-specific domain expansions, including B-box domain additions in HvCO clades. Tissue-specific expression patterns further supported functional divergence among paralogs. Notably, *VRN2*, a canonical floral repressor associated with winter growth, was retained in spring genotypes, challenging its presumed exclusive role in vernalization. Discrepancies between *VRN1* expression, *VRN2* deletion, and growth habits implicated additional regulatory mechanisms. These findings highlight the power of pangenomes in resolving gene family complexity, refining annotations, and advancing the understanding of CCT genes to enhance barley resilience and adaptability.

## Introduction

The CONSTANS (CO), CONSTANS-LIKE (COL), and TIMING OF CAB EXPRESSION1 (TOC1) or CCT motif genes are a highly conserved family of plant genes that regulate critical developmental and physiological processes, including flowering time, circadian rhythms, photoperiodic responses, and abiotic stress tolerance (Putterill et al., 1995; Cockram et al., 2007). These genes are characterized by a conserved CCT domain located at their C-terminus, which plays a role in DNA binding and protein-protein interactions (Shen et al., 2020). The CCT family is classified into three major clades based on their N-terminal domains: the CCT MOTIF FAMILY (CMF), COL, and PSEUDO-RESPONSE REGULATOR (PRR). COL proteins, such as CONSTANS (CO) in *Arabidopsis thaliana* and Heading date 1 (Hd1) in rice (*Oryza sativa*), contain one or two B-box-type zinc finger domains (Putterill et al., 1995; Yano et al., 2000). CMF proteins, including Grain Number, Plant Height, and Heading date 7 (Ghd 7) in rice (Weng et al., 2014), lack a conserved N-terminal domain. PRR proteins, such as TOC1/PRR1 in Arabidopsis and Ghd7.1 (OsPRR37) in rice, contain pseudo-receiver domains (Strayer et al., 2000; Murakami et al., 2005). CCT genes, distinguished by their evolutionarily conserved sequences and diverse functions, represent prime targets for improving yield and resilience in cereals (reviewed in Liu et al., 2020).

Recent studies in barley (*Hordeum vulgare*), a widely cultivated cereal crop, have revealed the role of CCT motif genes in regulating important agronomic traits. For example, *HvCMF4* is essential for floral development, and mutations in this gene cause primordia abortion and pollination failure by impairing rachis greening and plastidial energy supply (Huang et al., 2023). Other *CMF* genes, such as *HvCMF3* and *HvCMF7*, are crucial for chloroplast development with *HvCMF7* being important for chloroplast ribosome biogenesis and *HvCMF3* regulating chloroplast formation (Li et al., 2019; Li et al., 2021). Mutations in *HvCMF7* result in variegated leaves, while defects in *HvCMF3* lead to chlorosis and thylakoid abnormalities, highlighting their distinct but overlapping functions. *HvCO1* and *HvCO2*, barley’s closest homologs of *AtCO*, regulate flowering by activating *HvFT1* in response to photoperiod (Campoli et al., 2012a). *HvCO1* accelerates flowering under both long and short days, while *HvCO2* promotes flowering in a *Ppd-H1* (a PRR domain gene)-dependent manner (Turner et al., 2005; Campoli et al., 2012b). In winter barley, *HvFT1* activation is controlled by vernalization. Prior to vernalization, VERNALIZATION 2 (VRN2), a zinc-finger CCT motif protein (also known as ZCCT), acts as a floral repressor, preventing premature flowering (Yan et al., 2004b; Mulki and von Korff, 2016). After vernalization, *VRN1*, a MADS-box transcription factor, suppresses *VRN2* and activates *HvFT1*, promoting flowering through a positive feedback loop (Yan et al., 2006; Li et al., 2015). Phylogenetic and comparative analysis of CCT motif genes across species suggests they originated before the monocot/dicot divergence (Cockram et al., 2012). However, many *CMF* and *CO* genes within these families still have undefined functions, and the genetic diversity of these genes, especially in relation to global barley diversity, has yet to be fully explored.

Genome-wide characterization of gene families is a fundamental approach to understanding gene functions and their evolutionary dynamics. Early research, including the first characterization of barley *COL* genes, primarily relied on bacterial artificial chromosome (BAC) libraries (Yu et al., 2000; Griffiths et al., 2003; Yan et al., 2004b; Faure et al., 2007). With the increasing availability of complete genome sequences, these early methods have been greatly improved. For instance, the spring barley cultivar Morex was the first barley genome to be fully sequenced (Mascher et al., 2017) and subsequently better annotated (Monat et al., 2019). While these genomic resources enabled the development of a comprehensive barley gene family database (Li et al., 2022), the reliance on short-read sequencing for genome assembly has posed challenges in accurately resolving complex genomic regions (reviewed in Michael and VanBuren, 2020), such as the later-on detected structural variations (Jayakodi et al., 2020). Furthermore, single-genotype-based analyses are limited by the absence of key genes in the reference genotype. For example, *VRN2*, an essential regulator of vernalization and flowering time in winter barley, is absent from the Morex BAC library (Yan et al., 2004b). These limitations have hindered the full characterization of gene families and their functional diversity.

The recent development of a barley pangenome, which includes 76 diverse genotypes assembled using long-read HiFi sequencing, offers a transformative resource for genomic research (Jayakodi et al., 2024). This high-quality pangenome not only provides improved genome assemblies but also offers detailed gene annotations and orthologous relationships across genotypes. The pangenome incorporates genotypes with diverse phenotypic traits, such as growth habit (winter or spring), presenting a unique opportunity to explore the genetic basis of these traits. However, the link between these phenotypic differences and genetic variations in CCT motif family genes remains unclear, despite their known roles in plant architecture, flowering time, and vernalization regulation.

In this study, we adopt a pangenome perspective to classify CCT motif genes in barley, aiming to elucidate their genetic diversity, evolutionary patterns, and functional roles across diverse genotypes. By moving beyond traditional protein database-based analysis, we validated presence/absence variations (PAVs) and copy number variations (CNVs) directly from genome assemblies. This approach revealed significant genetic features, including potential reading frame variations in *HvCO1*, natural frameshift mutations in *HvCO10*, and copy number variations in *VRN2*. Notably, complete copies of *VRN2* were identified in multiple spring barley genotypes, challenging the long-standing assumption that the absence of *VRN2* is the sole genetic determinant of vernalization requirement.

## Results

### Identification of CCT motif genes in 76 barley pangenome assemblies

To systematically identify all CCT motif genes, we employed a conventional motif search approach using the barley pangenome version 2 (BPGv2) protein sequence database (panbarlex.ipk-gatersleben.de, Jayakodi et al., 2024). HvCO and HvPRR candidates were classified based on the overlap between identified CCT motif hits and corresponding B-box or PRR motif hits, respectively (Figure 1). In addition to the expected HvCMF and VRN2, the identified CCT motif-only candidates also include members of the GATA family of transcription factors, which are conserved in Arabidopsis and rice (Reyes et al., 2004). For nomenclature and validation of the motif-classified sets, we compared the identified sequences with previously published data (Griffiths et al., 2003; Campoli et al., 2012a; Cockram et al., 2012). We selected the winter barley cultivar Akashinriki as the reference for nomenclature, as some candidates, including VRN2 and HvPRR59, were already undetected in Morex at this stage (Dataset S1). Each candidate in Akashinriki was assigned a name according to its corresponding reference. Interestingly, contrary to the expectation of multiple copies (Dubcovsky et al., 2005), only a single copy of VRN2 was detected. Additionally, Akashinriki HvCO1 was recovered from a B-box motif-only candidate, suggesting a possible CCT motif deletion (Figure 1). Nevertheless, the phylogenetic relationships largely align with the findings in Morex (Huang et al., 2023), with the addition of the nomenclature for HvCMF2, HvCMF9, HvCMF12, HvCMF14. The assigned names were then propagated across all 76 genotypes based on established orthologous relationships (i.e., orthogroup ID, panbarlex.ipk-gatersleben.de, Jayakodi et al., 2024).

**Figure 1.**
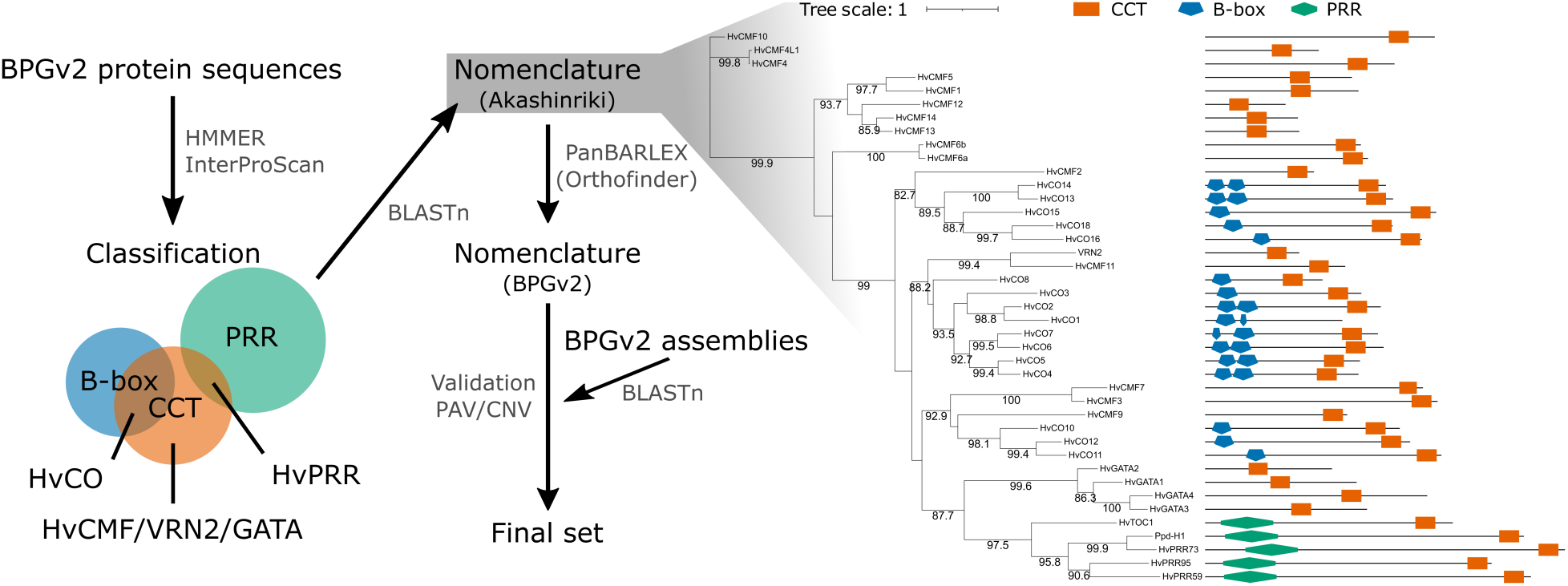
CCT motif proteins in barley and the identification pipeline. The identification process began with motif searches against the barley pangenome version 2 (BPGv2) protein sequences, followed by classification. Nomenclature was determined across all 76 genotypes based on orthologous relationships. Presence/absence variations (PAVs) and copy number variations (CNVs) were directly validated from the BPGv2 assemblies. The phylogenetic tree represents the identified candidates (prior to validation) in Akashinriki, including their expected domains, and was constructed using the maximum likelihood IQ-TREE JTT+F+I+G4 model with 1,000 bootstrap replications (values ≥ 80 are displayed).

The conservation of each candidate across the 76 genotypes allowed the classification of genes as core or dispensable within the pangenome context (Marroni et al., 2014). Of the 41 genes, 25 were present in all 76 genotypes and were considered as core genes, likely responsible for the major phenotypic traits (Figure 2A). The remaining genes were categorized as dispensable exhibiting greater diversity and potentially contributing to phenotypic plasticity. Thirteen of these dispensable genes were present in at least 72 genotypes, while *HvCO4*, *HvCO10*, and *VRN2* were found in only 69, 62, and 55 genotypes, respectively.

**Figure 2.**
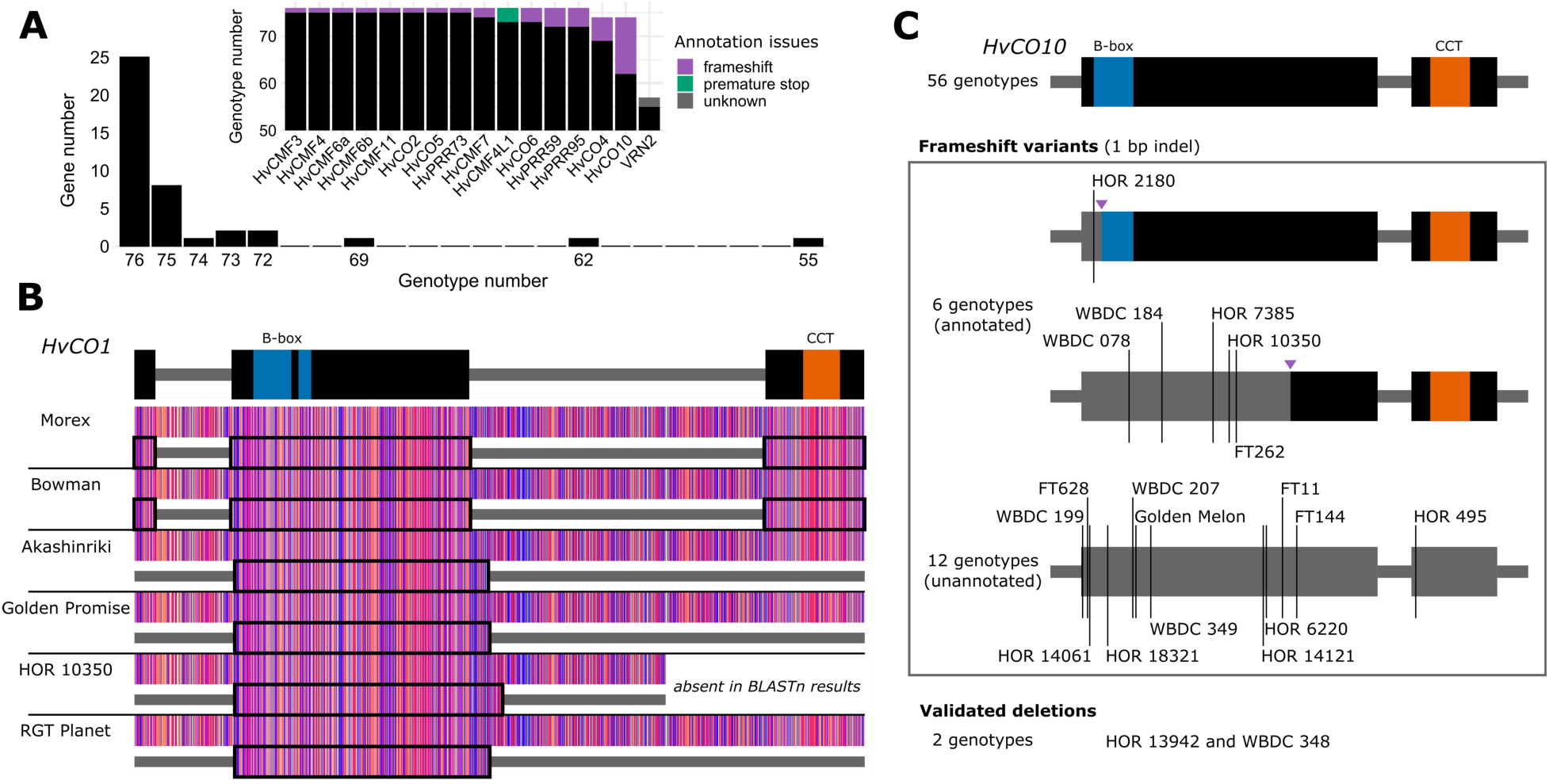
Validation of PAVs in barley CCT motif genes. (A) Presence/absence variations (PAVs) of CCT motif genes across 76 barley genotypes. Twenty-five genes were present in all 76 genotypes, while the PAVs of the remaining genes were validated and characterized, with potential annotation issues highlighted (inner plot). (B) Reading frame variations of annotated *HvCO1* across selected genotypes, leading to potential deletions in the CCT motif. Nucleotides are color-coded according to the ‘Zappo’ scheme in the sequence alignment. (C) Frameshift variants and deletions identified in *HvCO10*. Purple arrows indicate alternative translation start sites. Morex was used as a reference for the gene models in (B) and (C).

### Validation of PAVs and CNVs in barley CCT motif genes

Although protein databases and gene annotation-based methods are widely used for gene family identification, even in the pangenome era (Tong et al., 2025), careful validation of presence/absence variations (PAVs) and copy number variations (CNVs) remains crucial to prevent annotation errors, particularly when annotations rely on projections (Jayakodi et al., 2024). To verify the variations in so-far identified candidates, such as deletion of the CCT domain in Akashinriki HvCO1 and the absence of the expected VRN2 copies, we conducted a systematic search of all candidates using their genomic DNA sequences directly against the genome assemblies (Figure 1; Dataset S2).

While identified in all 76 genotypes, many core genes, including *HvCO1*, exhibited reading frame variations. For example, genotypes such as Golden Promise, HOR 10350, and RGT Planet, showed CCT domain deletions in HvCO1 at the protein level, similar to Akashinriki (Figure 2B). In these genotypes, the first and last exons of *HvCO1* were not transcribed and/or annotated compared to Morex and Bowman, leading to the loss of the CCT domain while retaining intact B-box domains. However, this may be due to annotation mistakes, as the genomic DNA sequences of *HvCO1* remain intact and highly conserved across these genotypes. The genotypes presented as examples here were all part of the first version of the barley pangenome (BPGv1), where gene annotations were primarily based on transcript evidence (Jayakodi et al., 2024; Guo et al., 2025). However, the annotated exons in both Morex and Akashinriki did not align with exons supported by RNA-seq and Iso-seq data (Figure S1), suggesting that the annotated coding regions may have been derived mainly from projections, potentially leading to inaccuracies. Similar variations in coding regions were observed in other CCT motif genes, including *HvCMF1*, *HvCMF3*, *HvCMF4L1*, *HvCMF10*, *HvCO2*, *HvCO18*, *HvGATA4*, and *HvPRR59* (Table S1). These findings highlight the importance of carefully validating gene annotations, with experimental confirmation needed to determine the accurate reading frame of these genes. Moreover, the extended open reading frame observed in HOR 10350 may result from frameshift insertions, while the deletion of the CCT domain could be validated by the absence of the corresponding coding regions. These variations in the predefined core genes indicate that an accurate definition of core genes should also consider the conservation of functional domains.

In addition to validating the identified candidates, our approach uncovered novel candidates that were undetectable at the protein level (Figure 2A, inner plot; Table S1). Many of these were absent due to frameshift mutations and/or premature stop codons, particularly in the case of *HvCO10*, where twelve genotypes lacked annotations and two exhibited true genomic deletions (Figure 2C). Furthermore, six genotypes with annotated *HvCO10* were also affected by frameshift mutations (1 bp insertion or deletion, indel). However, alternative translation start sites allowed them to remain in frame: in HOR 2180, this resulted in a partial deletion of the B-box domain, while in the other five genotypes, approximately half of the reading frame, including the CCT domain, was retained. Meanwhile, among the unannotated genotypes, HOR 14121, HOR 6220, FT11, FT144, and HOR 495 were predicted to contain only the B-box domain, highlighting *HvCO10* as one of the most diverse CCT genes with extensive natural mutations. This diversity suggests that *HvCO10* is undergoing active evolutionary diversification, potentially driven by selective pressures such as environmental adaptation or functional redundancy. In contrast, similar natural variants were identified in only a few genotypes of other CCT motif genes (Figure 2A; Table S1). However, for genes with known functions, such as *HvCMF3*, *HvCMF4*, *HvCMF7*, and *HvPRRs*, these newly discovered natural variants provide valuable resources for further functional studies.

We then investigate genes expected to have additional paralogs. Previous phylogenetic analysis indicated that barley (Morex) contains *HvCMF4* paralogs, *HvCMF4L1* (also known as *HvCMF8*) and *HvCMF4L2*, which have partially redundant roles in spikelet development regulation (Huang et al., 2023). While only HvCMF4 and HvCMF4L1 were detected at the protein level using motif searches (Figure 1; Figure 2A), we successfully detected *HvCMF4L2* and two additional copies by searching with *HvCMF4L1* genomic DNA sequences. To validate the phylogenetic relationships of these newly identified copies, we constructed a phylogenetic tree including all CCT motif proteins, excluding frameshift variants (Figure 3A). As expected, the new copies clustered together with HvCMF4 in the same clade and were designated as HvCMF4L3 and HvCMF4L4. Sequence analysis revealed that HvCMF4L2 and HvCMF4L3 exhibited partial deletions of the CCT domain (Figure 3B), which explained why they were not initially detected through motif searches. In contrast, HvCMF4L4 lacked the entire CCT domain and shared only partial sequence conservation with HvCMF4, HvCMF4L2 and HvCMF4L3, excluding it from being classified as a CCT motif protein. HvCMF4L2 was identified in only 18 genotypes, while the newly discovered HvCMF4L3 was more abundant, present in 54 genotypes. Both genes were dispensable and located on chromosome 3H at different positions, suggesting that their presence was not due to intrachromosomal translocation, as supported by limited sequence conservation and the co-occurrence of both paralogs in certain genotypes, such as 10TJ18. Consistent with previous findings (Huang et al., 2023), these results indicate that HvCMF4L1, HvCMF4L2, and HvCMF4L3 likely arose from independent duplication events of HvCMF4. However, the functional role of HvCMF4L3 remains to be determined.

**Figure 3.**
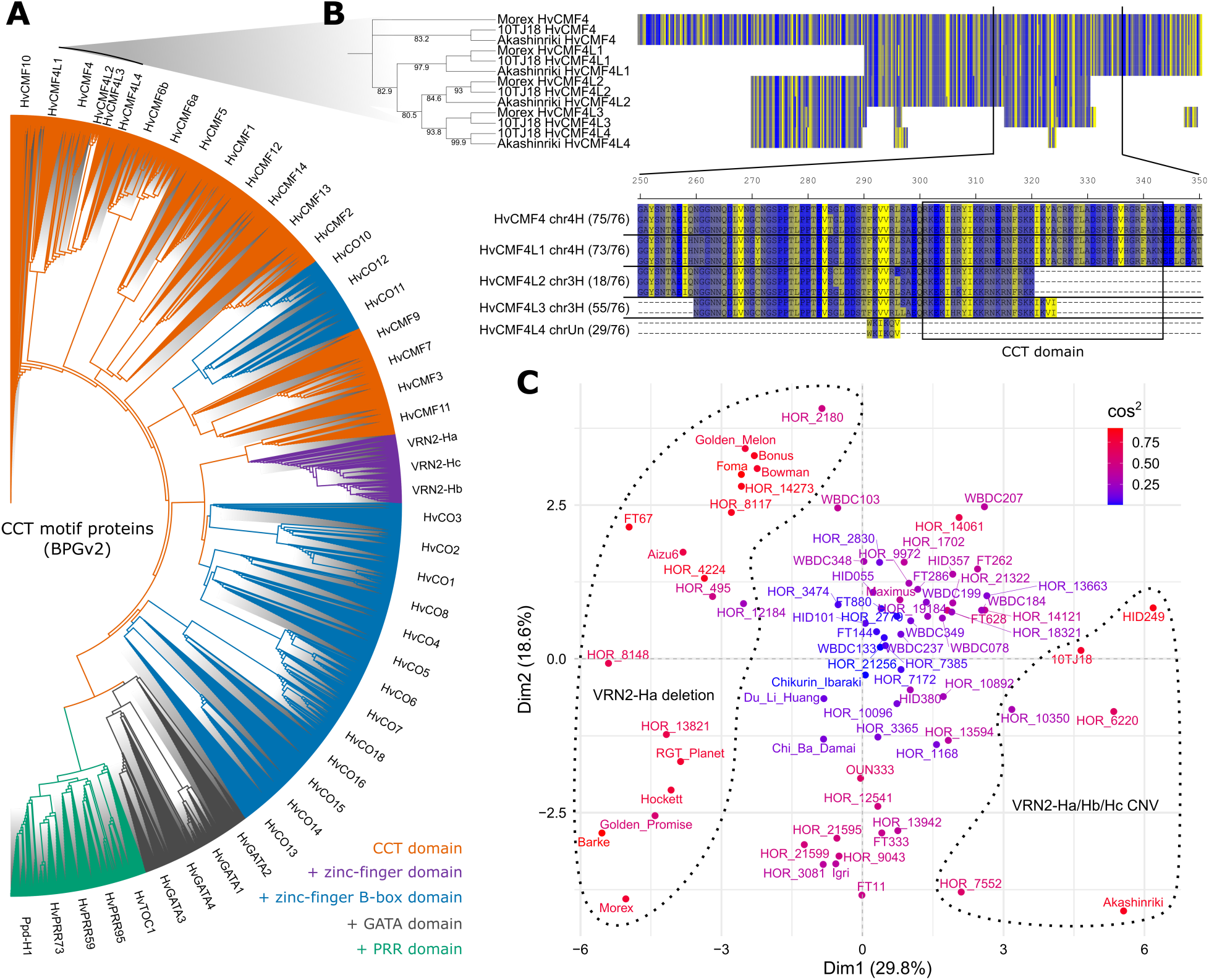
Diversity of CCT motif proteins in the barley pangenome. (A) Phylogeny of CCT motif proteins across 76 genotypes. The phylogenetic tree was constructed using the maximum likelihood IQ-TREE JTT+F+I+G4 model. Branches are colored based on the presence of CCT domain alone (except for HvCMFL4) or the addition of other domains. (B) Phylogeny of HvCMF4 and HvCMF4L in selected genotypes. The phylogenetic tree was constructed using IQ-TREE JTT+I model with 1,000 bootstrap replications (values ≥ 80 are displayed). Amino acids are color-coded according to the ‘Strand’ scheme in the sequence alignment. The CCT domain is based on Morex HvCMF4. The number in brackets next to the sequence alignment indicates the number of genotypes containing the corresponding annotated copy. (C) Principal component analysis (PCA) of CCT motif protein variation across the barley pangenome. PCA was based on all CCT motif proteins as shown in (A), grouped by genotype. Cosine squared (cos^2^) values reflect the contribution of each genotype to the principal components. The genotype clusters, marked by dashed lines, are based VRN2-Ha deletion and VRN2-Ha/Hb/Hc copy number variations, manually assigned according to Table S2.

As the most ‘dispensable’ CCT motif gene present in the ‘shell pangenome’ of barley, *VRN2* is expected to exhibit multiple copies, referred to as *VRN2* (or *ZCCT*)*-Ha*/*Hb*/*Hc* (Dubcovsky et al., 2005; von Zitzewitz et al., 2005). These copies encode proteins containing a CCT domain, with VRN2-Ha and VRN2-Hb also possessing an additional zinc-finger domain. While only a single copy of VRN2 was annotated in the barley pangenome (Figure 1), additional copies were identified by searching genome assemblies using published reference sequences (Yan et al., 2004b). In total, 57 genotypes were found to contain at least one *VRN2* copy (Figure 2A; Table S2). However, as these copies were not supported by pan-transcriptome data (Figure S2), their reading frames were predicted and subsequently classified and validated through phylogenetic analysis (Figure 3A).

### Refined phylogeny of CCT motif proteins in barley

With the inclusion of the missing copies, we were able to establish a comprehensive phylogenetic overview of CCT motif proteins at the barley pangenome level (Figure 3A). Despite observed genotypic variations (e.g., partial deletions, Figure 2B, C), each CCT member distinctly grouped into separate clades, reinforcing their correct nomenclature based on orthologous relationships.

Consistent with single-genotype phylogenies based on MorexV2 (Huang et al., 2023) and Akashinriki (Figure 1), the pangenome phylogeny revealed that HvCMF10, HvCMF4, and HvCMF4L were the most ancient forms of the CCT family. They were followed by the highly similar and neighboring copies HvCMF6a and HvCMF6b, whose diversification appeared to have occurred later—after HvCMF5, HvCMF1, HvCMF12, HvCMF14, and HvCMF13—when considering only Akashinriki (Figure 1). This underscored the advantage of using a pangenome approach for such phylogenetic analysis, as it minimized biases introduced by genotypic variations. The recent pan-transcriptome dataset based on BPGv1 provided additional insights into gene function through co-expression and tissue-specific expression patterns (Guo et al., 2025). Although *HvCMF4* and *HvCMF4L* were largely absent in the dataset (Figure S2), we observed strong expression of *HvCMF6a* and *HvCMF6b* in caryopsis, as well as *HvCMF6a* in inflorescence, suggesting their roles in inflorescence development and grain filling, potentially mirroring the functions of *HvCMF4* and *HvCMF4L* (Huang et al., 2023).

In barley, the first B-box domain addition within the CCT family coincided with the divergence of two distinct clades: one consisting of HvCO12, HvCO11, and HvCO10, and the other containing HvCMF9, HvCMF3, and HvCMF7 (Figure 3A). Notably, only the first clade acquired an additional B-box domain, suggesting a potential, yet unknown, functional divergence driven by domain expansion. Interestingly, all three HvCOs in this clade contained only a single B-box domain and were phylogenetically distant from other HvCOs, some of which possessed two B-box domains. This suggests that the B-box domain addition likely occurred through independent evolutionary events. The CCT domain primarily facilitates DNA binding, while zinc-finger B-box domains are thought to play a crucial role in protein-protein interactions (Ben-Naim et al., 2006). Consistent with their functional conservation (Li et al., 2019; Li et al., 2021), HvCMF3 and HvCMF7 clustered within the same subclade, with *HvCMF3* showing high expression in shoots, while *HvCMF7* was broadly expressed across all five tissues (Figure S2; Dataset S3). However, it remains uncertain whether their role in chloroplast development emerged before or after their divergence from HvCMF9. If it evolved beforehand, it raises the question of how the addition of B-box domain might have influenced this function.

Similarly, the addition of the zinc-finger domain in VRN2-Ha was an independent evolutionary event that occurred during its diversification from HvCMF11 (Figure 3A). However, VRN2-Hb and VRN2-Hc appeared to have arisen through duplications of VRN2-Ha, with VRN2-Hc subsequently losing the zinc-finger domain. This was supported by the observations that all genotypes carrying VRN2-Ha also contained VRN2-Hb and VRN2-Hc, except for the wild barley (*Hordeum spontaneum*) genotype FT67, which only possessed VRN2-Hb and VRN2-Hc (Table S2).

Most of the remaining HvCOs have unknown functions, except for HvCO1 and HvCO2, which are conserved across species for their role in regulating flowering time (Campoli et al., 2012a). While both displayed significant genotypic variations due to complex splicing patterns (Figure 2B; Table S1), they each contained an additional B-box domain compared to HvCO3, from which they originated (Figure 1; Figure 3A). This indicated that the domain duplication may have contributed to their specific functions, distinguishing them from other HvCOs.

The phylogenetically most recent CCT members were those that had acquired zinc-finger GATA or PRR domains. GATA transcription factors, a group of DNA binding proteins found widely in eukaryotes (Lowry and Atchley, 2000), were represented by four members in barley corresponding to Arabidopsis class C GATA factors, which were characterized by the presence of both a TIFY and a CCT motif, although their specific functions remained unclear (reviewed in Schwechheimer et al., 2022). Transcriptome data indicated that *HvGATA2* was predominantly expressed in roots, *HvGATA1* was specific to inflorescences, while *HvGATA3* and *HvGATA4* exhibited overall higher expression levels particularly in inflorescences (Figure S2; Dataset S3). However, these members were likely dispensable within the CCT motif protein family and did not originate from HvCMFs. The addition of the CCT domain appeared to have supplemented their GATA-family related functions.

A similar hypothesis could be applied to the members with PRR domains; however, they represented a more complete family, including HvTOC1 (HvPRR1), HvPRR95, HvPRR59, HvPRR73, and Ppd-H1 (HvPRR37) (Campoli et al., 2012b). A modeling approach indicated that most of them were part of the barley circadian network, conserved in Arabidopsis (Müller et al., 2020). However, Ppd-H1 likely acquired its circadian functions after diverging into AtPRR3 and AtPRR7 in Arabidopsis (Turner et al., 2005). In barley, evident from its high expression levels in shoots and inflorescences (Figure S2; Dataset S3), *Ppd-H1* maintained its essential role in photoperiod response and flowering time regulation within a pathway that includes HvCOs (Campoli et al., 2012b). This suggested that the function of barley Ppd-H1 predates its later role as a core component of the circadian network in Arabidopsis. Additionally, HvTOC1 exhibited high expression in roots, indicating its potential involvement in root-specific circadian regulation.

### Barley pangenome insights into *VRN2* and growth habit variation

The refined phylogenetic analysis of CCT motif proteins in the barley pangenome revealed distinct clades, domain expansions, and potential functional divergences. However, the impact of genotypic variation remained unclear, as all genotypes clustered within the same clade for each CCT motif protein, despite the variations described above. To address this, we performed principal component analysis (PCA) on the same set of sequences (all CCT proteins) separated by genotypes. The PCA plot revealed that most genotypes, regardless of row-type or geographic origin, clustered together with lower cos^2^ values (purple and blue), indicating minimal contribution to variation (Figure 3C). In contrast, genotypes with stronger contributions (red) were primarily separated by Dim1 (29.8%), driven largely by PAVs and CNVs of VRN2 (Table S2), consistent with *VRN2* being identified as the most dispensable CCT motif gene in barley (Figure 2A).

To determine whether this PCA-based approach could reliably identify the most variant genes and to examine the role of Dim2 (18.6%) in separating the VRN2-Ha deletion cluster, we repeated the analysis excluding all VRN2-related sequences. Notably, genotypes previously separated by Dim2 were now separated along Dim1 (29.2%) in the new PCA plot (Figure S3A). To further explore this separation, we selected two sets of highly contrasting genotypes—Morex and Barke versus HOR 2180 and Bowman—and constructed a phylogenetic tree (Figure S3B). This analysis pinpointed HvCMF1 as the driver of the observed separation, which aligned with the presence of two major reading frame variants in HvCMF1 based on annotation (Figure S3C; Table S1). However, it is important to note that this could also be influenced by potential annotation errors, as previously observed in HvCO1 (Figure S1), necessitating further validation.

While most genotypes with VRN2 deletions were well-characterized spring barley cultivars (e.g., Morex, Barke, and Golden Promise), winter barley genotypes such as Aizu6, HOR 4224, and HOR 12184 also exhibited complete VRN2 deletions. Surprisingly, several spring barley genotypes retained intact VRN2-Ha/Hb/Hc copies, with HOR 6220 and HOR 7552 even carrying additional copies. These findings suggest that the relationship between spring growth habits and VRN2 deletions should be reconsidered. Since all identified *VRN2* copies were located in close proximity on chromosome 4H, we selected representative genotypes to validate CNVs and local chromosomal structures. Although both Igri and Akashinriki are winter barley cultivars, Akashinriki carried two additional *VRN2-Ha* copies, and an extra copy of *VRN2-Hb* and *VRN2-Hc* (Figure 3C; Table S2), likely resulting from regional duplication and/or translocation (Figure 4A).

**Figure 4.**
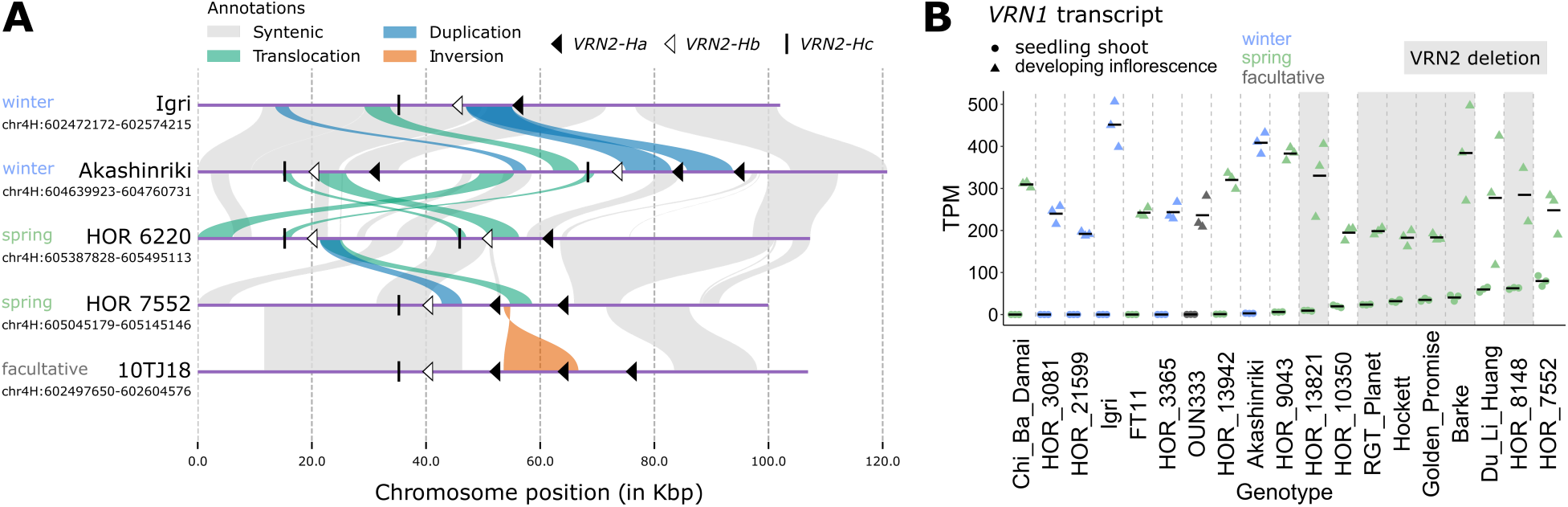
The presence of *VRN2* is not associated with winter growth habit in barley. (A) Synteny of genomic regions containing *VRN2* copy number variations in selected genotypes: including winter barleys Akashinriki and Igri, spring barleys HOR 6220 and HOR 7552, and the facultative barley genotype 10TJ18. *VRN2* copies are indicated based on their coordinates, located near regions with complex duplications, translocations, and inversion. (B) *VRN1* transcript levels in seedling shoots and developing inflorescences across 19 barley genotypes. Genotypes shaded in grey have deletions of all *VRN2* copies. Growth habit information in (A) and (B) is based on (Jayakodi et al., 2024).

Compared to Akashinriki, the spring barley landrace HOR 6220 contained only one *VRN2-Ha* copy, similar to Akashinriki’s second copy, while *VRN-Hb* and *VRN2-Hc* regions underwent complex translocations and duplications. Even between spring barley genotypes, *VRN2* CNVs were not entirely syntenic between HOR 6220 and HOR 7552.

In addition to winter and spring growth habits, some barley genotypes exhibit a facultative growth habit, characterized by spring-like growth without vernalization sensitivity but strong low-temperature tolerance (Von Zitzewitz et al., 2011). Similar to spring and winter barleys, facultative barleys in the pangenome displayed *VRN2* PAVs and CNVs (Figure 3C; Table S2). For instance, the additional *VRN2-Ha* copy in 10TJ18, compared to HOR 7552, likely resulted from local inversion and insertion (Figure 4A). As these copies are highly conserved, their identification was only possible through long-read sequencing-based genome assemblies. We confirmed that the entire *VRN2*-containing region in Akashinriki, which had the highest copy number, was supported by coverage within a single contig (contig_362, Jayakodi et al., 2024). Although synteny relationships varied in pairwise comparisons depending on the reference genotype, these findings highlight that *VRN2* PAVs and CNVs in barley are linked to complex genome rearrangements.

Although no experimental data currently support the regulatory roles of the newly identified *VRN2* PAVs and CNVs in defining growth habits, pan-transcriptome resources (Guo et al., 2025) may provide insights into the vernalization transcriptional hub *VRN1* across diverse genotypes. *VRN1* transcript levels from seedling shoots before vernalization and developing inflorescences after vernalization offer valuable information on vernalization requirements and effects. As expected, vernalization increased *VRN1* transcript levels from minimal in all winter barley genotypes, confirming their vernalization sensitivity (Figure 4B). In contrast, most spring barleys exhibited high *VRN1* transcript levels before vernalization, indicating no vernalization requirement. However, exceptions were observed in some spring or facultative genotypes, such as Chi Ba Damai, FT11, OUN333, HOR 13942, and HOR 9043, which showed expression patterns similar to winter barleys. This may be explained by the presence of potentially functional *VRN2* copies in these genotypes (Figure 3C; Table S2). This remains unvalidated, as *VRN2* transcripts were absent from the current PanBaRT20 database (Guo et al., 2025), which likely resulted in their annotation omission. Interestingly, other *VRN2*-containing spring genotypes, such as Du Li Huang and HOR 7552, exhibited high *VRN1* expression, raising questions about the exact repression role of *VRN2* in barley flowering regulation. Taken together, our results highlight the need for a systematic pangenome-level investigation of vernalization regulatory mechanisms in barley.

## Discussion

Advancements in plant pangenomics have revolutionized the study of gene families, offering unprecedented opportunities to explore genetic diversity, evolutionary dynamics, and functional variation within and across species (Tao et al., 2019; Jayakodi et al., 2025). However, transitioning from single-reference genome analyses to pangenome-based frameworks necessitates careful methodological considerations to ensure accurate and comprehensive gene family characterization. Our study highlights the importance of integrating multiple data sources and validation strategies to overcome the limitations of traditional approaches, particularly in structurally complex regions and highly variable gene families such as barley CCT motif genes.

Accurate detection of presence/absence variations (PAVs) and copy number variations (CNVs) is pivotal in pangenome-based gene family analysis. The examination of CCT motif proteins within the barley pangenome has revealed notable gene annotation discrepancies, especially concerning core genes like *HvCO1*. These discrepancies suggest that the observed variations may result from annotation inaccuracies rather than genuine genetic differences. Current gene annotation strategies, including those employed in the barley pangenome (Jayakodi et al., 2024), rely on both transcript evidence and projection methods. While pan-transcriptome data offer genotype-specific evidence across five tissues (Guo et al., 2025), they are limited by factors such as sampling at a single time point, which may lead to missing information in the transcriptome data of CCT genes, thereby shifting annotations towards projections. A significant portion of the barley transcriptome (∼84%) is considered rhythmic and is regulated by the circadian clock under long-day conditions (Müller et al., 2020). For instance, the major flowering regulator *HvCO1* exhibits expression patterns predominantly at the end of the day and in the evening, in contrast to genes such as *Ppd-H1* (Campoli et al., 2012a; Zahn et al., 2023). These findings highlight the need for careful validation of gene annotations, integrating comprehensive transcriptomic and genomic data to accurately define core genes and their functional domains.

While motif-based searches can identify core CCT motif genes, direct examination of genome assemblies has uncovered potentially functionally disruptive frameshift mutations, such as those in *HvCO10*. These findings emphasize that relying solely on protein-level annotations can overlook structural variants, especially for genes under positive selection pressures that exhibit high diversity within species. Conversely, such novel natural alleles serve as valuable resources for understanding gene functions. For instance, the diverse functions of the barley flowering time regulator *EARLY FLOWERING 3* have been elucidated through natural allelic variation (Zahn et al., 2023; Zhu et al., 2023; Huang et al., 2024). Determining whether these variations contribute to proteome complexity necessitates large-scale mass spectrometry-based proteomics analyses (Tress et al., 2017).

Long-read sequencing technologies have been instrumental in resolving intricate genomic architectures and uncovering structural variations that often elude detection in short-read assemblies (Michael and VanBuren, 2020). For example, the identification of *VRN2* copies in spring barley genotypes challenges previous assumptions that vernalization requirements are exclusively governed by *VRN2* deletions (Yan et al., 2004b; Landis et al., 2024). The potential involvement of other repressors, such as the MADS-box transcription factor gene *ODDSOC2* (Greenup et al., 2010; Dixon et al., 2019; Jacott and Boden, 2020), in barley’s vernalization response warrants further investigation. Unlike Arabidopsis, where vernalization is mediated through *FLOWERING LOCUS C* (*FLC*), temperate grasses such as wheat and barley utilize a distinct pathway centered on *VRN1* (reviewed in Xu and Chong, 2018). However, transcript-level analyses of *VRN1* across 19 genotypes revealed no clear correlation between its expression and growth habits or *VRN2* deletions. A comprehensive understanding of this regulation requires examination of genetic variations in *VRN1*. Targeted sequencing of *VRN1* in 1,000 barley accessions from diverse geographical regions revealed ten alleles containing intron deletions or insertions that modulate its pre-vernalization expression independently of cold exposure (Hemming et al., 2009). Natural deletions in the *VRN1* promoter region (recessive *vrn1* alleles) correlate with growth habits in wheat but only partially explain vernalization differences in barley (Yan et al., 2004a; von Zitzewitz et al., 2005). Barley’s diploid genome, with fewer vernalization gene copies, compared to polyploid wheat, may render its flowering regulation more sensitive to genetic variation.

Recent wheat pangenome studies demonstrate that *VRN1* variations in subgenome A alone determine growth habits (Jiao et al., 2024), underscoring the need for similar investigations in the barley pangenome to unravel species-specific regulatory mechanisms. The observed discrepancies between *VRN1* transcript levels and growth habits may also result from the nuanced nature of growth habit classifications. Traditionally, growth habits are defined based on vernalization requirements and/or sensitivity, which are quantitative traits often approximated by measuring the days to flowering in non-vernalized plants. This approach may not align strictly with a binary winter or spring classification. For instance, genotypes with complete or partial deletions of *VRN2* can still exhibit varying degrees of vernalization sensitivity, intermixed with genotypes containing intact *VRN2* copies (Muñoz-Amatriaín et al., 2020).

In addition to genetic factors, the ‘winter memory’ mechanism underlying vernalization is epigenetically regulated. In wheat and barley, the upregulation of *VRN1* during vernalization is associated with a shift in histone modification marks, transitioning from the repressive H3K27me3 to the activating H3K4me3 (Oliver et al., 2009; Diallo et al., 2012). Advanced epigenomic profiling techniques have mapped chromatin states of key vernalization genes both before and after vernalization, uncovering vernalization-related regulatory elements, including distal accessible chromatin regions upstream of *FT* (Liu et al., 2024). The identification of these regulatory elements, which requires functional genomic annotations (i.e., by epigenomic profiling), were not covered in the current analysis of CCT genes.

The current set of barley pangenome genotypes, featuring high-quality assemblies and verified *VRN2* variations, offers an excellent starting point to revisit and refine the definitions of growth habits and vernalization mechanisms in barley.

## Methods

### Identification of CCT motif genes in the barley pangenome

All proteins containing CCT (Pfam ID: PF06203), B-box (PF00643), and PRR (PF00072) domains were identified using HMMER/3.1b2 (Finn et al., 2011) based on the barley pangenome version 2 (BPGv2) protein sequence database (panbarlex.ipk-gatersleben.de; Jayakodi et al., 2024). Overlapping domain sets were used to classify candidates into HvCMFs (CCT only), HvCOs (CCT with B-box), and HvPRRs (CCT with PRR) (Dataset S1). Proteins with potential CCT domain deletions but retaining B-box or PRR domains were also retrieved from the corresponding HMMER results. To validate HMMER-based classification, InterProScan/5.57 (Jones et al., 2014) was conducted using the MorexV3 protein database (Jayakodi et al., 2024), targeting CCT (IPR010402), B-box (IPR000315), and PRR (IPR017053) domains. A few candidates, including functionally annotated *TIFY* genes and *FAR1-related sequence 5*, were initially identified as containing potential CCT domains but were excluded due to high E-values (> 1).

Gene nomenclature was first assigned candidates identified in winter barley cultivar Akashinriki using BLASTp/2.13.0 (Camacho et al., 2009) against published sequences (Griffiths et al., 2003; Campoli et al., 2012a; Cockram et al., 2012). Since the phylogenetic relationships of barley GATA candidates were unclear compared to Arabidopsis and rice (Reyes et al., 2004), they were sequentially named as HvGATA1 (orthogroup ID: N0.HOG0052967), HvGATA2 (N0.HOG004833), HvGATA3 (N0.HOG0057350), and HvGATA4 (N0.HOG0060472). These names were then assigned to corresponding candidates in the other 75 genotypes based on orthologous relationships (Jayakodi et al., 2024). At this stage, the candidates remained highly conserved across genotypes, as orthologous relationships were established using OrthoFinder (Emms and Kelly, 2019).

To verify presence/absence variations (PAVs) and copy number variations (CNVs), genomic coordinates for each gene were extracted based on annotations (Table S1, Dataset S2) and were used for BLASTn/2.13.0 (Camacho et al., 2009) searches across all 76 genome assemblies (Jayakodi et al., 2024). The resulting genomic DNA sequences were retrieved using SAMtools/1.16.1 (Li et al., 2009) and aligned with annotation-based coding sequences from all 76 genotypes. Sequence alignments were performed using MUSCLE (Edgar, 2004) in AliView (Larsson, 2014) and visualized with the R package ggmsa (Zhou et al., 2022). For newly identified sequences or copies, gene names and orthogroups were assigned based on genomic coordinates if annotations were available. Since *VRN2-Hb* and *VRN2-Hc* were absent from annotations in all 76 genotypes, all *VRN2*-related targets were classified according to their reference sequences: AY485977.1 (*VRN2-Ha*), AY485978.1 (*VRN2-Hb*), and AY687931.1 (*VRN2-Hc*) (Yan et al., 2004b; Dubcovsky et al., 2005). The open reading frames of *VRN2-Hb* and *VRN2-Hc* were predicted using AUGUSTUS web interface (Stanke et al., 2006).

### Phylogenetic analysis

While validating PAVs and CNVs, novel frameshift and/or premature stop variants were identified but excluded from phylogenetic analysis to prevent potential biases. However, newly identified copies of HvCMF4L and VRN2 were included. Sequence alignments were conducted using MAFFT/7.490 (Katoh and Standley, 2013), and maximum likelihood phylogenetic analysis was performed with IQ-TREE/2.2.2.6 (Minh et al., 2020), employing 1,000 bootstrap replications. The best-fit amino acid substitution model was selected based on the Bayesian Information Criterion (BIC) before constructing the phylogenetic tree. Phylogenetic trees were then visualized and annotated using iTOL (Letunic and Bork, 2007).

### PCA

To analyze sequence variations among genotypes, PCA was performed using k-mer frequency-based representations of amino acid sequences. The sequences were grouped by genotype to construct a k-mer (*k* =3) frequency matrix. Biostrings (Pages et al., 2013) was used to read and process amino acid sequences, extracting 3-mer sub-sequences for each genotype and counting their occurrences. The union of all k-mers across genotypes was compiled to define the matrix columns. To account for partial deletions, PAVs, and CNVs, each row in the matrix, representing a genotype, was normalized by the total k-mer count for that genotype. PCA was then performed using the FactoMineR package (Lê et al., 2008) with the normalized k-mer frequency matrix as input. The PCA scores were extracted, and individual genotypes were visualized in two-dimensional space, with points colored according to their cosine squared (cos^2^) values, indicating their contributions to the principal components. Visualization was conducted using the factoextra package (Kassambara and Mundt, 2017), applying a blue-to-red gradient to represent the degree of contribution. Labels were adjusted to minimize overlap, ensuring clear depiction of clustering patterns.

### Analysis of *VRN2* regional synteny

In the selected genotypes Igri, Akashinriki, HOR 7552, HOR 6620, and 10TJ18, genomic regions containing all identified *VRN2* copies were extracted from their respective genome assemblies using SAMtools/1.16.1 (Li et al., 2009) based on the coordinates. A phylogenetic tree was first generated constructed (as described above) to determine the order of pairwise sequence comparisons. Pairwise alignments were performed using minimap2/2.24 (Li, 2018) with the ‘-ax sr’ setting to align short genomic regions between genotypes. The alignments were processed with SyRI/1.6 (Goel et al., 2019) to retain alignment information, and structural variations were identified for each pairwise comparison. Plotsr/0.5.2 (Goel and Schneeberger, 2022) was used to visualize the synteny relationships, with a window size of 1,000 bp, and a distance threshold of 1,000 bp. *VRN2* gene positions were annotated according to their genomic coordinates.

### Analysis of the pan-transcriptome

Transcriptome data, quantified as transcripts per million (TPM), for all five tissues were obtained from the barley genotype-specific reference transcript datasets (RTDs, Guo et al., 2025). Due to inconsistencies between pan-transcriptome gene IDs and pangenome gene (or orthogroup) IDs, gene IDs were linked based on genomic coordinates. Genes aligned with multiple truncated transcripts were excluded from the dataset. Transcripts of all CCT genes were extracted from 20 PBGv1 genotypes (Dataset S3), including *HvCO1* transcripts that aligned with two additional neighboring genes (Figure S1). Average transcript levels were calculated based on biological replicates (n = 2-3) and visualized using the ComplexHeatmap package (Gu et al., 2016). *VRN1* transcripts were extracted from shoot and inflorescence tissues and visualized using the R package ggplot2 (Wickham et al., 2016). Morex *VRN1* (*HORVU.MOREX.PROJ.5HG00452020*) transcripts were absent from the datasets.

To examine the transcriptome data used for gene annotation in *HvCO1*, genotype-specific RTDs, derived from RNA-seq and Iso-seq, along with exon annotations, were extracted from Morex and Akashinriki (Guo et al., 2025) and visualized using IGV (Robinson et al., 2011).

## Acknowledgements

This work was supported by the German Ministry of Education and Research (BMBF) (grant number FKZ 031B1224).

## Supplementary data

**Table S1.** Genomic validation of CCT genes.

**Table S2.** PAVs and CNVs of *VRN2* in the barley pangenome.

**Dataset S1.** Barley CCT motif proteins identified through HMMER search.

**Dataset S2.** Barley CCT genes identified by BLASTn.

**Dataset S3.** Transcriptome data of CCT genes across 20 barley genotypes.

## Figure legends

**Figure S1.** Absence of transcriptome support for *HvCO1* annotation. Genomic regions containing *HvCO1* (grey shaded area) in Morex and Akashinriki. Gene annotations tracks (Jayakodi et al., 2024) are shown alongside genotype-specific reference transcript datasets (RTDs) derived from RNA-seq and Iso-seq, as well as exon annotations (Guo et al., 2025).

**Figure S2.** Expression variability of CCT genes across 20 barley genotypes. The heatmap illustrates the average transcript levels (n = 2-3), quantified in transcripts per million (TPM, Guo et al., 2025) for all CCT genes across five tissues. For each tissue, data from 20 genotypes are displayed in the following order (left to right): Akashinriki, Barke, Chi Ba Damai, Du Li Huang, FT11, Golden Promise, Hockett, HOR 10350, HOR 13821, HOR 13942, HOR 21599, HOR 3081, HOR 3365, HOR 7552, HOR 8148, HOR 9043, Igri, Morex, OUN333, and RGT Planet. The gene order is determined by the phylogenetic tree (Figure 3A). *HvCO1* transcripts, indicated by an asterisk, were aligned with two additional neighboring genes (Figure S1).

**Figure S3.** HvCMF1 variation in the barley pangenome. (A) Principal component analysis (PCA) of CCT motif protein variation across the barley pangenome, excluding VRN2. Cosine squared (cos^2^) values reflect the contribution of each genotype to the principal components. Arrows highlight selected contrasting genotypes: Morex and Barke versus HOR 2180 and Bowman. (B) Phylogeny of CCT motif proteins, excluding VRN2, in selected genotypes. The phylogenetic tree was constructed using the maximum likelihood IQ-TREE JTT+F+I+G4 model, with branches colored according to genotype groups, emphasizing HvCMF1 as the most divergent protein between groups. (C) Two major HvCMF1 variants identified in the barley pangenome, potentially explaining the genotypic separation observed in (A). Amino acids are color-coded according to the ‘Strand’ scheme in the sequence alignment.

