## Supplementary material for "Pangenome insights into structural variation and functional diversification of barley CCT motif genes": Figure S1

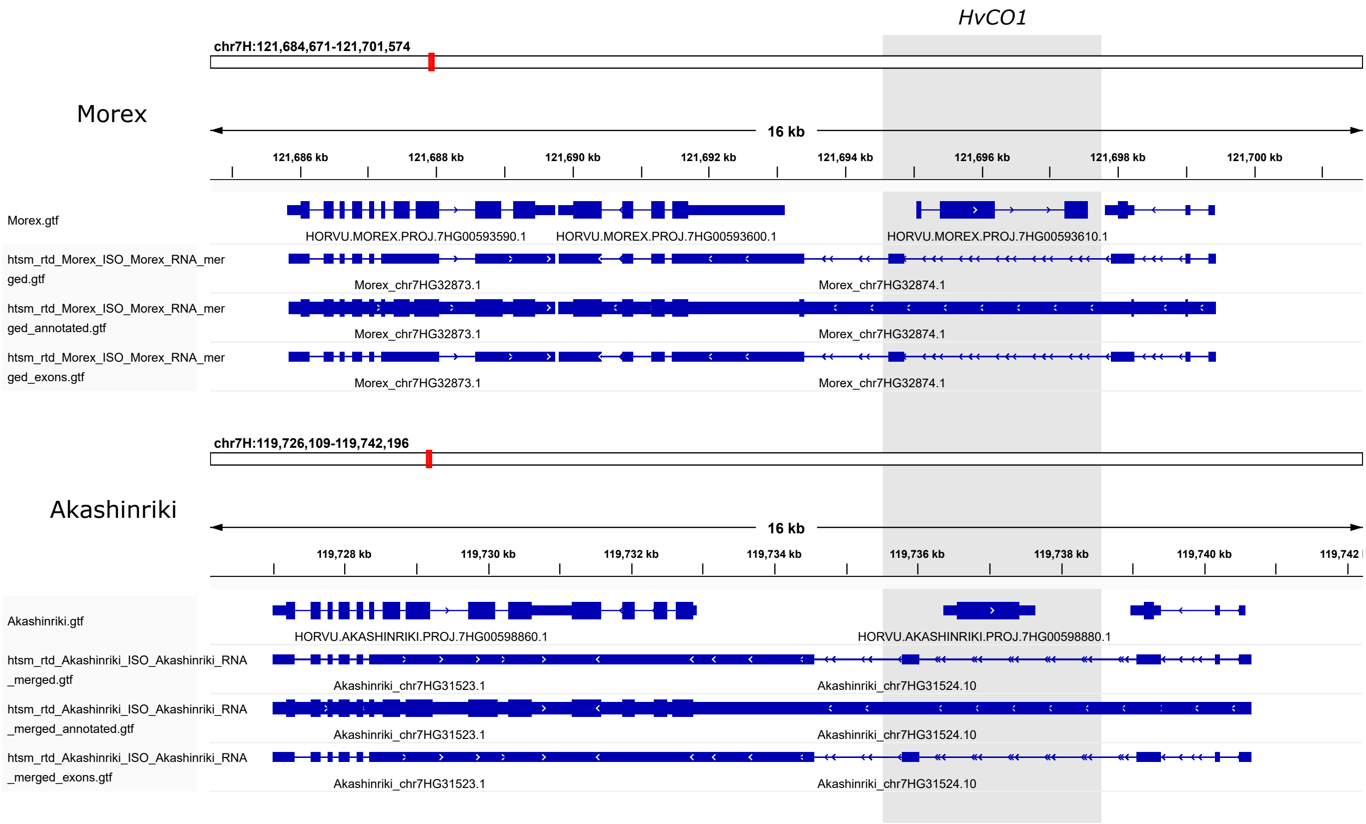


**Figure S1.** Absence of transcriptome support for *HvCO1* annotation. Genomic regions containing *HvCO1* (grey shaded area) in Morex and Akashinriki. Gene annotations tracks (Jayakodi et al., 2024) are shown alongside genotype-specific reference transcript datasets (RTDs) derived from RNA-seq and Iso-seq, as well as exon annotations (Guo et al., 2025).
