## Supplementary material for "Pangenome insights into structural variation and functional diversification of barley CCT motif genes": Figure S2

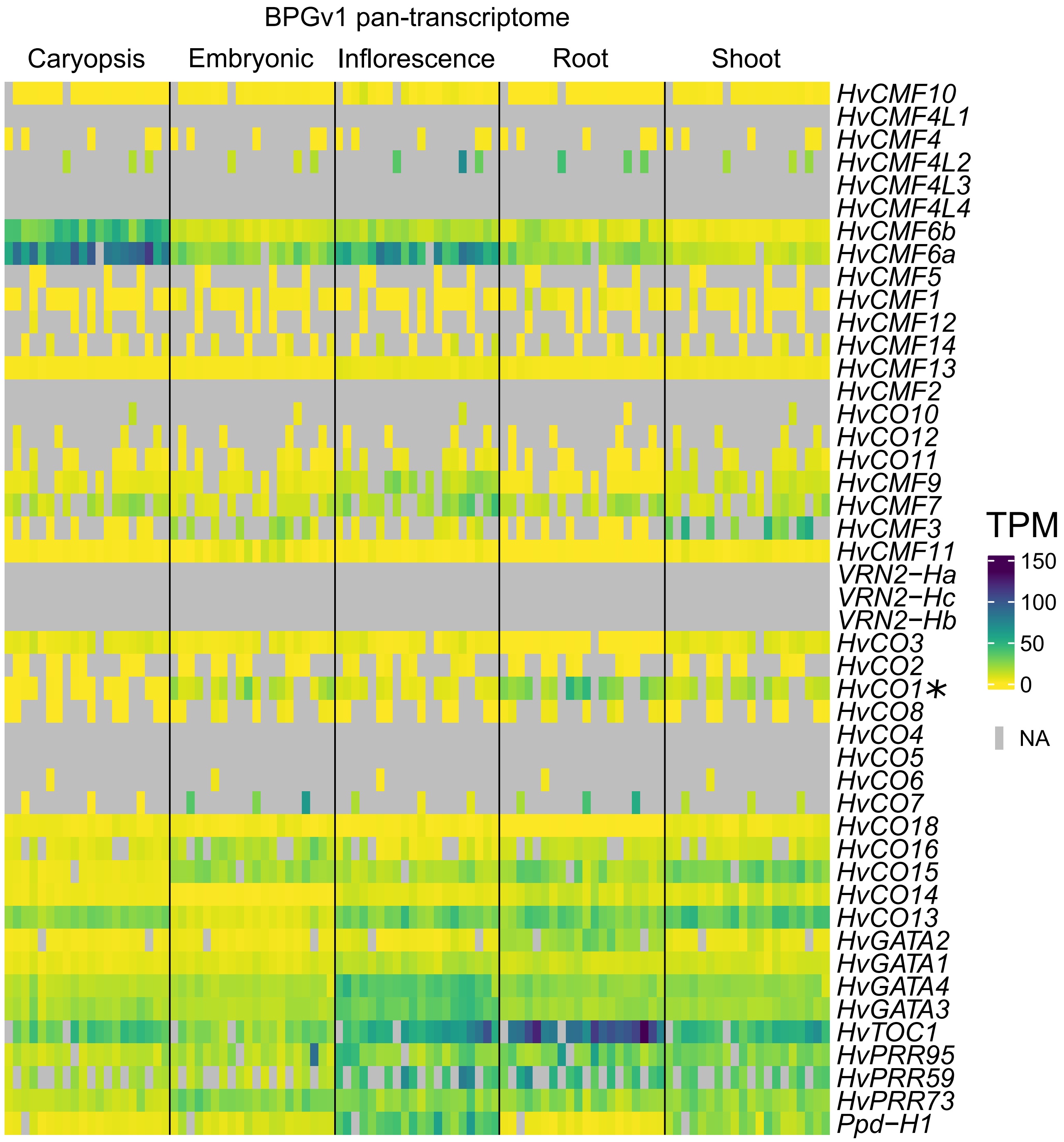


**Figure S2.** Expression variability of CCT genes across 20 barley genotypes. The heatmap illustrates the average transcript levels (n= 2-3), quantified in transcripts per million (TPM, Guo et al., 2025) for all CCT genes across five tissues. For each tissue, data from 20 genotypes are displayed in the following order (left to right): Akashinriki, Barke, Chi Ba Damai, Du Li Huang, FT11, Golden Promise, Hockett, HOR 10350, HOR 13821, HOR 13942, HOR 21599, HOR 3081, HOR 3365, HOR 7552, HOR 8148, HOR 9043, Igri, Morex, OUN333, and RGT Planet. The gene order is determined by the phylogenetic tree (Figure 3A). *HvCO1* transcripts, indicated by an asterisk, were aligned with two additional neighboring genes (Figure S1).
