## Supplementary material for "Pangenome insights into structural variation and functional diversification of barley CCT motif genes": Figure S3

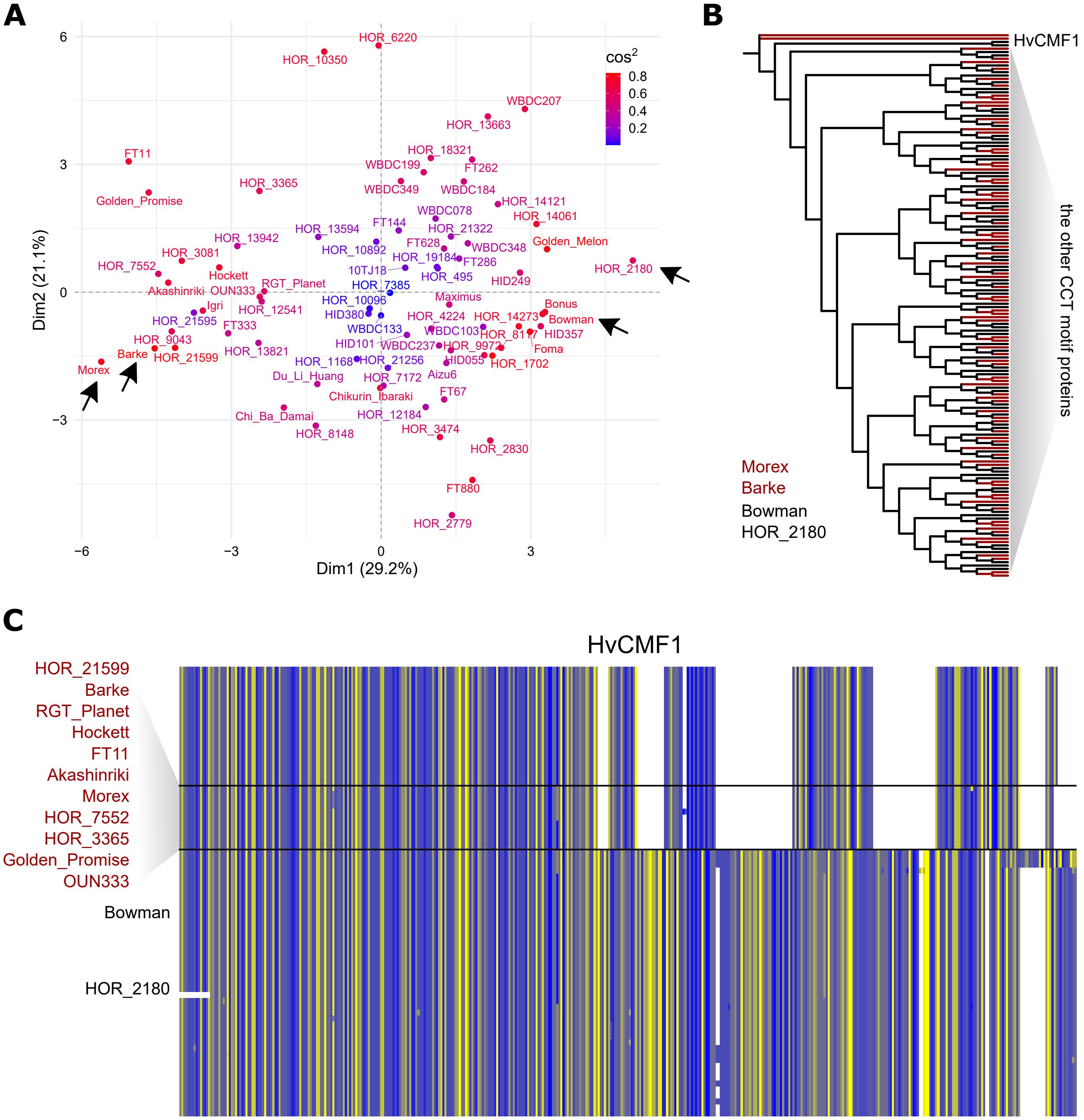


**Figure S3.** HvCMF1 variation in the barley pangenome. (A) Principal component analysis (PCA) of CCT motif protein variation across the barley pangenome, excluding VRN2. Cosine squared ($cos^2$) values reflect the contribution of each genotype to the principal components. Arrows highlight selected contrasting genotypes: Morex and Barke versus HOR 2180 and Bowman. (B) Phylogeny of CCT motif proteins, excluding VRN2, in selected genotypes. The phylogenetic tree was constructed using the maximum likelihood IQ-TREE JTT+F+I+G4 model, with branches colored according to genotype groups, emphasizing HvCMF1 as the most divergent protein between groups. (C) Two major HvCMF1 variants identified in the barley pangenome, potentially explaining the genotypic separation observed in (A). Amino acids are color-coded according to the ‘Strand’ scheme in the sequence alignment.
